# Combination of dynamic turbidimetry and tube agglutination to identify procoagulant genes by transposon mutagenesis in *Staphylococcus aureus*

**DOI:** 10.1101/426783

**Authors:** Dong Luo, Qiang Chen, Bei Jiang, Shirong Lin, Linfeng Peng, Lingbing Zeng, Xiaomei Hu, Kaisen Chen

## Abstract

Agglutinating function is responsible for an important pathogenic pattern in *S.aureus.* Although the mechanism of aggregation has been widely studied since *S.aureus* has been found, the agglutinating detailed process remains largely unknown. Here, we screened a transposon mutant library of Newman strain using tube agglutination and dynamic turbidmetry test and identified 8 genes whose insertion mutations lead to a decrease in plasma agglomerate ability. These partial candidate genes were further confirmed by gene knockout and gene complement as well as RT-PCR techniques. these insertion mutants, including NWMN_0166, NWMN_0674, NWMN_0756, NWMN_0952, NWMN_1282, NWMN_1228, NWMN_1345 and NWMN_1319, which mapped into coagulase, clumping factor A, oxidative phosphorylation, energy metabolism, protein synthesis and regulatory system, suggesting that these genes may play an important role in aggregating ability. The newly constructed knockout strains of *coa, cydA* and their complemented strains were also tested aggregating ability. The result of plasma agglutination was consistent between *coa* knockout strain and *coa* mutant strain, meanwhile, *cydA* complement strain didn’t restored its function. Further studies need to confirm these results. These findings provide novel insights into the mechanisms of aggregating ability and offer new targets for development of drugs in *S.aureus.*

## INTRODUCTION

*Staphylococcus aureus* (*S.aureus*) is still an important clinical pathogen, which can lead to localized infections such as skin and soft issue infection and systemic severe infections, such as septicemia and sepsis ^[1,2]^. Since first discovered in England, methicillin-resistant *S. aureus* (MRSA) has quickly been a predominant epidemiological pathogen worldwide ^[3]^. Because of higher drug-resistance rate, MRSA is more difficult to treat than methicillin-susceptible *S. aureus* (MSSA). However, the effective clearance of *S. aureus* depends on not only drug-resistance but also comprehensive spectrum of virulence factors ^[4,5]^.

As an important pathogen that widely distributes in the environment, *S. aureus* can cause infections in almost all body sites, and infectious ability is highly associated with the comprehensive effect of virulence factors ^[6]^. These virulence factors include hemolysin, staphylococcal protein A (SPA), Panton-valentine leukocidin (PVL), coagulase, enterotoxin, and so on. Different virulence factors have different mechanism in pathogenicity, including enhanced adhesion, strengthening invasion, immunosuppression, phagocytosis inhibition, dissolving neutrophils and so on ^[7,8]^. As a result, vaccines targeting the conservative amino acid sequences of virulence factor can protect *S. aureus* infections ^[9,10]^.

It is still important to know the pathogenic mechanisms of different virulence factors in order to effectively prevent *S. aureus* infections. In brief, there are two pathogenic mechanisms for these virulence factors. First, partial virulence factors, including SPA, DNase, FnbB, Lipase and SasX, facilitate the bacteria adhere to host cells and destroy their local barrier ^[11]^. Secondly, other virulence factors including ClfA, FnbpA, Coa, and all enterotoxins antagonize or escape host immune responses. The mechanical barrier in *S. aureus* is an important immune escape pattern, by secreting agglutination factor, plasma and blood cells agglutination is activated and thus forming fibrous sheet structure, which functions as a mechanical barrier and wraps *S. aureus* in abscess, avoiding attack of immune cells ^[12]^. McAdow and his colleagues ^[10]^ confirmed that *S. aureus* is easy to be removed by immune cells when the aggregation ability depress, even it is not easy to form bacteremia.

The first step for mechanical barrier formation in *S. aureus* is to form fibrous net-like structure by promoting plasma or blood agglutination. Generally speaking, the formation of staphylococcal abscess needs the local blood cells and plasma to agglutinate first, and then congeal blood by making fibrinogen transform fibrin to shape fibrin mesh. In this process, coagulase activation plays a crucial role. Although coagulase can’t catalyze the conversion of prothrombin into thrombin, it can alter the three-dimensional structure of prothrombin, expose the enzyme active center and execute thrombin function. *S. aureus* secretes two kinds of coagulases, free coagulase and bound coagulase (clumping factor). The former can promote agglutination of blood cells and plasma, forming the inner core of coated vesicles ^[13]^.The latter forms the outer fiber network and cleaves alpha and beta chain of fibrinogen ^[14]^. In addition, *S. aureus* can combine with specific sites of soluble fibrin under the intermediary of ClfA to form complexes, leading neutrophils, macrophages and other immune killing cells fail to contact *S. aureus*, thus preventing *S. aureus* from being killed and cleared ^[15]^. It is confirmed that the ability of abscess formation is limited when *S. aureus* lacks free coagulase or von-willebrand factor-binding protein(vWbp), and the ability is further restricted when both of them lack ^[16]^. Studies have shown that the symptoms of invasive arthritis relieve when *S. aureus* lacks *clfA* gene^[17]^, but no effect was observed when ClfA antibody was used for treatment of staphylococcal sepsis ^[18]^. It is suggested that some unknown genes involved in the process of plasma coagulation except the widely-known *coa, vwbp* and *clfA* genes. In addition, recent studies have also confirmed that thickening of bacterial cell wall can reduce the ability of plasma agglutination of *S. aureus*, probably due to inhibiting the exocrinosity of plasma coagulase ^[19]^.

To further understand the mechanisms of plasma agglutination, we constructed and subsequently screened a transposon mutant library of *S. aureus* Newman strain, and identified 8 genes by using combination of tube aggregation and dynamic turbidmetry tests. We selected coa and cydA gene for further functional study by gene knockout and complementation technique. The results proved that these genes shall have important link with plasma agglutination.

## MATERIALS AND METHODS

### Bacterial Strains, Plasmids, Antibiotics, and Growth Conditions

*E. coli* strains DH5α and DC10B, *S. aureus* strain RN4220 were used in DNA cloning. *S. aureus* Newman strain was used as a parent strain for screening. The plasmids pID408, pRB473, and pKOR1 were used for construction of transposon library, genetic complementation, and homologous recombination, respectively ^[20]^. Antibiotics were used at the following concentrations, unless otherwise indicated: erythromycin, 20μg/ml; ampicillin, 100μg/ml; chloramphenicol, 20μg/ml. Typtic soy agar (TSA) and typtic soy broth (TSB) were used for *E.coli* and *S. aureus* growth.

### Construction of transposon mutant library

*S. aureus* transposon mutant library was constructed based on previous description. At first, *S. aureus* RN4220 and Newman strain were prepared from log phase cultures grown in TSB broth (Oxiod company), then centrifuged precipitation by 5,000 r/min to collect bacteria, and competent cell was acquired by 3 times sucrose washing. Secondly, the temperature-sensitive plasmid pID408 containing transposon Tn917 was successively transformed into *S. aureus* RN4220 and Newman competent cells by electroporation (diameter of electrode cup =0.2cm, voltage=2.5kV, resistance=100Ω, capacitance=25μF) using GenePulser Xcell^TM^ (Bio-Rad). After resuscitation, the cells were spread on TSA plates containing 20μg/ml erythromycin and 20μg/ml chloramphenicol and incubated at 30°C. Then the Newman strain containing pID408 plasmid was cultured at 43°C and 250rpm for 18h to induce transposon insertion. The induced clones were collected and incubated on TSA plates with 20μg/ml erythromycin and with or without 20μg/ml chloramphenicol. The transposon mutant would not survive on plates containing chloramphenicol for missing of pID408. Transposon mutagenesis bias was tested by inverse PCR. Finally, about 10,000 mutant clones were picked and cultured in TSB broth and then stored as transposon mutant library at −80°C.

### Tube agglutinating test (TAT)

Mutant *S.aureus* Newman was grown onto 5% sheep blood plates at 37°C for 24h. Each single colony was turned to TSB (10ml) for culture by 250r/min at 37°C for 24h, respectively. Then *S.aureus* culture was centrifuged 6,000g for 30min. After discarding the supernatant fluid, the sediment was resuspended in sterile normal saline, and the bacterial concentration being 4.0 McFarland units.

TAT was performed with 50μl bacterial suspension and 450μl mixed citrate plasma from healthy volunteers (no obvious hemolysis and jaundice) in plastic tube (13mm diameter). After gently mixing, the tubes were incubated at 37°C. Clotting was evaluated hourly in the first 4 h and then at 16 h timepoint. An opaque clot which remained in place when the tube was tilted 90° from the vertical was identified as a positive result.

### Dynamic turbidimetry test (DTT)

Bacteria suspension was prepared as described above. The DTT was performed in 96-well plate with 10μl bacteria suspension and 90μl mixed citrate plasma from healthy volunteers (no obvious hemolysis and jaundice). The reactive system was monitored at 37°C, and OD_340_ values were measured. Agglutinating status was evaluated every 2 minutes during 6 h incubation. Positive DTT result was identified when OD_340_ value was more than 0.02 compared with the initial.

### Identification of mutated genes

The insertion site of the transposon was confirmed by the resistance marker detection^[21]^. First, the overnight culture of each mutant with reduced coagulating ability in TSB was lysed in lysis buffer, subsequently extracted bacterial DNA by alkaline lysis. Secondly, *EcoR*I enzyme digested the genome at 37°C for 12h, then T4 DNA ligase linked target fragment and transformed into *E.coli* DH5α by chemical transfection method. Screened and collected self-connected fragment of plasmid with TSA plates containing 20μg/ml AMP. Collected plasmid and used and design primer (5’-AGAGAGATGTCACCGTCAAGT-3’) at inverted repeat region of the near end of the erythromycin resistance gene of Tn917. After digestion with self replicating genes and source of resistance to starting point, *S.aureus* genomic DNA sequence in a new plasmid; according to the sequencing result with standard strains of *S.aureus* in Http://blast.ncbi.nlm.nil.gocv/-Blast.cgi were compared, obtained by Genebank transposon insertion sites for specific gene analysis and forecast the possible function.

### Gene knockout and complementation

The temperature-sensitive shuttle plasmid pKOR1 was used to knockout the candidate genes (*coa* and *cydA*) associated with decreased ability of agglutination by homologous recombination as described previously ^[20]^. Taking *cydA* as an example, approximately 1kb upstream and downstream sequences of *cydA* were amplified (primers used: *cydA*-UF, *cydA*-UR; *cyd A*-DF, *cydA*-DR) and cloned to plasmid pKOR1. The recombinant knockout plasmid was transformed into DC10B by electroporation, in which the plasmid was modified, and then into *S. aureus* Newman by electroporation. Homologous recombination was carried out by a two-step procedure. In the first step, under the selective pressure of high temperature (42°C) and chloramphenicol (10μg/ml), the plasmid was integrated into the chromosome of *S. aureus* through a single-crossover event. The integrated strain was then subcultured and the double-crossover occurred subsequently in the absence of antibiotic selection, thus acquiring either wild-type strain or the *cydA* knockout mutant depending upon where the recombination event occurred. In the second step, the culture was steaked on the plates containing 1μg/ml anhydrotetracycline to select the *cydA* knockout mutant strains, the strain carrying the plasmid sequence of the chromosome could not survive. Ten colonies were picked, and the correct knockout strain was verified by PCR and gene sequencing. The candidate gene and its upstream promoters were PCR-amplified using primers cydA-IF and cydA-IR. The restriction endonuclease of *BamH*I and *Kpn*I were used to cleavage plasmid pRB473 and PCR-amplified product. After transformation, DC10B was modified and the recombinant plasmids were transformed into the cydA deletion mutants via electroporation.

### Real-time qPCR

RT-qPCR was used to assess the expression levels of *cydA* and *coa* in Newman wild type, knockout, and complemented strains. Culture of *S. aureus* strain Newman grew overnight in TSB at 37°C for 8 hours. The cultures were centrifuged and washed twice with PBS. RNA isolation was performed according to manufacturer’s instruction (Sangon Biotech Co., Ltd., Shanghai). Primers were designed using Primer Express software (Version 2.0, Applied Biosystem). The reverse transcription was performed using Super-Script III First-Strand Synthesis (Takara Bio). RT-qPCR was performed using SYBR Premix Ex Taq II (Takara Bio). The amplicon of 16S rRNA was used as an internal control. Cycling conditions were 95°C for 30s and followed by 40 cycles of 95°C for 5s, 60°C for 30s. Relative expression levels were determined by the comparative threshold cycle (ΔΔCt) method.

## RESULTS

### Construction of mutant library and screening of clones with reduced plasma agglutination capacity

Mutant library was constructed by random mutation of Newman standard strain using Tn917 transposon, and then these clones whose agglutination ability disappeared or decreased were screened by dynamic turbidimetry and tube agglutination technique. A total of about 1,000 clones were collected, and 82 clones showed decreased or disappeared agglutination ability.

## Identification of insertion genes and bioinformatics analysis

In order to determine the inactivation gene inserted into the transposon in each mutant strain, the insertion site of each transposon was identified by the resistance rescue marker method, and the nucleotide sequence downstream of the insertion site of *S.aureus* was obtained by sequencing. The insertion site of the transposon in the genome of *S.aureus* was confirmed by comparison with the whole genome sequence of Newman wild strain, and the predicted gene number of the corresponding insertion mutation was obtained by analysis. The insertion sites of 82 strains were identified, and 76 mutants with reduced or disappeared agglutination capacity were identified. Six strains were not identified, which may be caused by the poor quality of DNA template extraction or the long restriction fragment near the insertion site, which leads to the formation of plasmids too large to be transformed successfully.

The identified mutant genes and their possible functions were searched and analyzed according to the information in NCBI. A total of 8 predicted functional genes were involved in 76 mutant strains identified (Table 2). Among them, 12 mutants transposons were inserted into NWMN_0674; 19 mutant transposons were inserted into the predicted gene NWMN_0952; All 7 mutant transposons were inserted into the predicted gene NWMN_0166; All 5 mutant transposons were inserted into the predicted gene NWMN_0756; three mutant transposons were inserted into the predicted gene NWMN_1345; 13 mutant transposons were inserted into the predicted gene NWMN_1282; 16 mutant transposons were inserted into the predicted gene NWMN_1319; one mutant transposons was inserted into the predicted gene NWMN_1228.

**Table 1.**
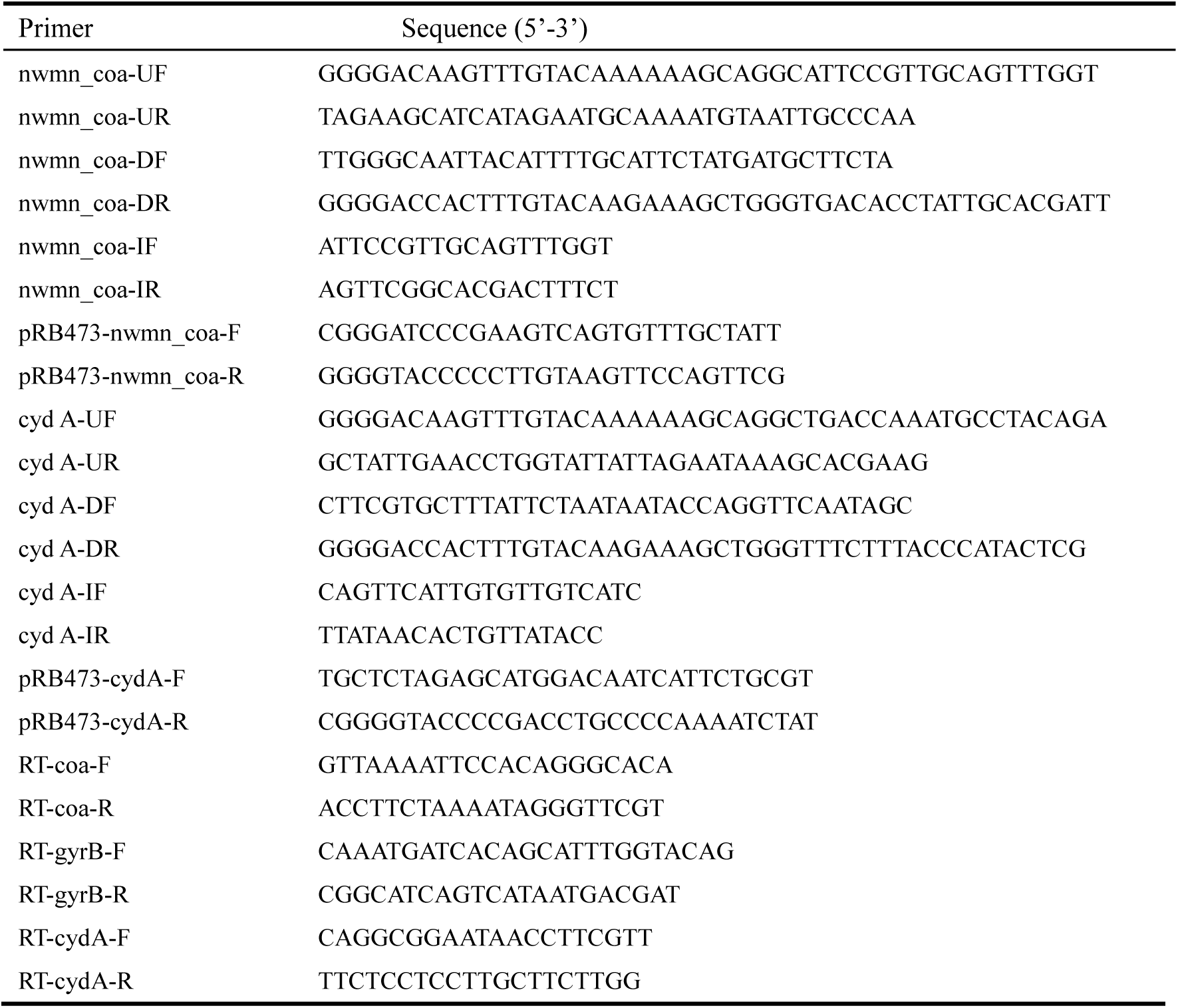
Primers used in this study.

**Table 2.** The biological information of the mutated genes.

| Quantity | Agglutinant ability | Located Gene | Predictive function |
| --- | --- | --- | --- |
| 12 | No agglutination | NWMN_0674 | SaeRS system |
| 3 | Decreased<br>agglutination | NWMN_1345 | Hypertonic permeable resistance protein Ebh |
| 5 | No agglutination | NWMN_0756 | Clumping factor A |
| 7 | No agglutination | NWMN_0166 | Coagulase |
| 19 | No agglutination | NWMN_0952 | Ubiquinone cytochrome reductase subunit I |
| 13 | Decreased<br>agglutination | NWMN_1282 | Indole-3-glycerol synthase |
| 1 | Decreased<br>agglutination | NWMN_1228 | Threonine |
| 16 | No agglutination | NWMN_1319 | Functional unknown protein |

**Table 3.**
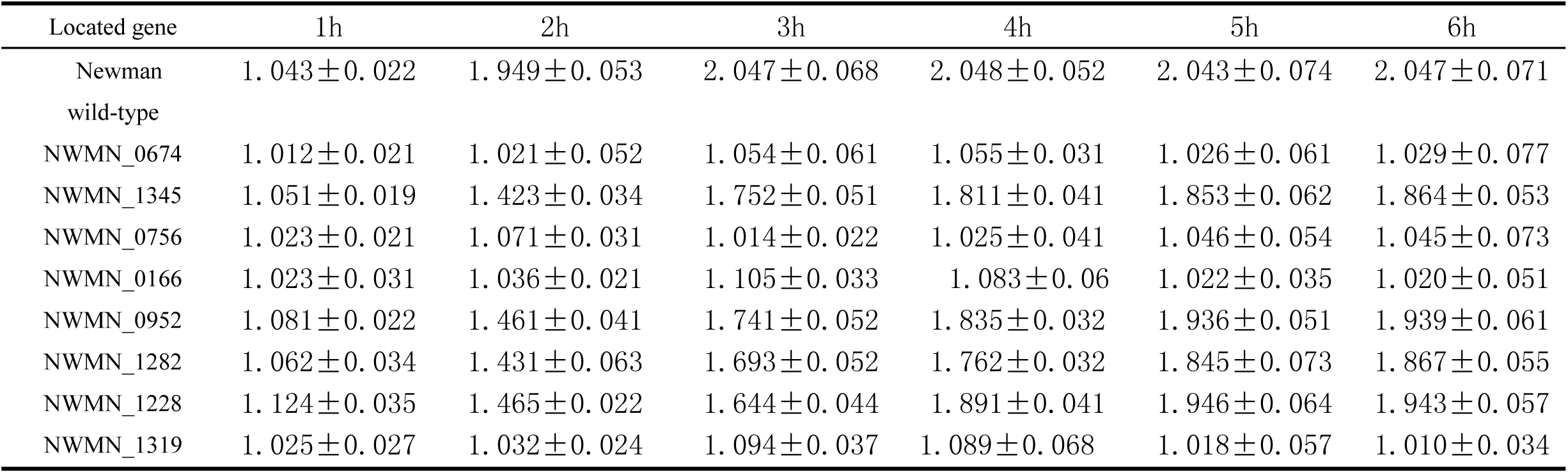
OD_340_ Values of dynamic turbidimetry of different mutant strians.

Among the inserting mutations identified, there were 12 mutants of the predictive and global regulatory system, involving one gene. A total of 15 strains of exocrine proteins were involved in 3 genes. And 19 energy metabolism mutant strains, involving 1 genes; There are 14 mutants related to protein synthesis and 2 genes involved. The other 16 mutants were located in unknown functional protein.

## Validation of screened genes

Two genes were selected for validation, one of which was specifically associated with plasma agglutination of *S.aureus* as the *coa* gene encoding plasma coagulase; The another gene was screened from this mutant library with a relatively high frequency, and no literature has been reported at present.

*coa* gene is recognized as the main gene to promote plasma agglutination, and the result of this experiment is consistent with those reports in many previous literatures ^[22]^. The mutant strain couldn’t promote plasma agglutination, and the corresponding complementary strain completely restored the physiological phenotype of wild strains. We didn't get the same result as we expected for *cydA* gene. However, we confirmed that the gene has been expressed by using fluorescence quantitative PCR as well as corresponding *coa* gene. The reason is that *cyd* gene is a complex encoded by three genes, although the *cydA* subunit is normally expressed, the other 2 subunits have not performed their function. Because the nucleotide encoded by three genes is larger than 10KB and can’t be transferred to plasmid at the same time, our current method can’t be further studied (the results weren’t shown).

## DISCUSSION

In this study, we identified 8 functional genes that could be associated with promoting plasma agglutination in *S. aureus*. The inactivation by insertion mutation led to decreased plasma agglutination. Because of the difficulty and large amount of work involved in validating all 8 genes, in this study we focused on validating the *coa* and *cydA* genes by constructing new knockout mutants, which were identified in multiple insertion mutants, suggesting a strong possibility that these genes are involved in promoting plasma agglutination in *S. aureus* Newman. Another reason to construct new knockout mutants is to rule out the polar effect of transposon insertion on upstream and (or) downstream genes. The finding that the freshly constructed *coa,* and *cydA* knockout mutants also had the same dereasing plasma agglutination as the transposon mutants demonstrates that these genes are indeed involved in promoting plasma agglutination. Furthermore, complementation of *coa* could completely restore the agglutinating ability, but complementation of *cydA* didn’t restore the agglutinating ability. The reason is *cyd* gene is a complex which includes three genes, *cydA, cydB* and *cydX*, expression of these genes may be different in genome vs. plasmid-based complementation. Additionally, the influence of other genes in the plasmid and possible compensation by other genes in the genome may also be the reasons for partially restoration. Further studies are needed to validate other insertion mutants by gene knockout in future studies.

To the best of our knowledge, this is the first comprehensive study using a transposon mutant library screen to systematically identify promoting plasma agglutinating genes in *S. aureus*. Our study provides important insights into the mechanisms of promoting plasma agglutination in *S. aureus* and identifies new drug targets for virulence-targeted antibiotic development for the improved treatment of system infections.

The gene *cydA* encodes ubiquinone cytochrome oxidase subunit I, which is involved in bacterial electron transfer chain, and there are no studies in *S.aureus* that electron transfer are involved in coagulase production, although energy production genes still affect virulence factors expression ^[23,24]^. Our studies proved that SaeRS system, *ClfA, Coa* and *Ebh* genes are essential in plasma agglutination in *S.aureus* ^[25-27]^, and our research had same results. Additionally, our results showed that some protein synthesis genes, such as NWMN_1282, NWMN_1228 and NWMN_1277 mutants could decrease agglutination, the reason want to further elucidate.

Our previous study found that the gene coverage of the transposon mutant library was well when we randomly examined the inserted sites. However, when reviewing the literature, we found that some genes can also affect plasma agglutination, for example, agr system, arlSR system, rot and *vwbp* ^[28,29]^. It was regrettable that we couldn’t screen these genes in our study. The reasons might be the number of genes we screened and identified is too small. Future studies with more clones or with different mutant libraries may be needed to more comprehensively understanding of *S. aureus* decreased agglutinating genes.

In summary, we identified 8 genes, including 4 novel genes, which may associated with decreased agglutinating genes in *S. aureus.* These genes are involved in several pathways including protein synthesis, electron transport chain, DNA repair, genes regulation, and unknown functional genes. These findings provide novel insights into the mechanisms of agglutinating formation in *S. aureus,* and offer new targets for developing new antibiotics and vaccine that prevent and treat *S.aureus* infections.

## Author Contributions

KSC and XMH designed the work and revised the manuscript; DL, QC, BJ, SR L, and LFP completed all the experiments; DL, and ZLB performed the statistically analysis and made the figures; DL, and QC wrote the manuscript.

## Conflict of Interest Statement

The authors declare that the research was conducted in the absence of any commercial or financial relationships that could be construed as a potential conflict of interest.

## Acknowledgments

This work was supported by the key project funding of Jiangxi Provincial Education Department (GJJ160025) and Natural Science Foundation of Jiangxi Province (20171BAB205074).

---

**Figure 1.**
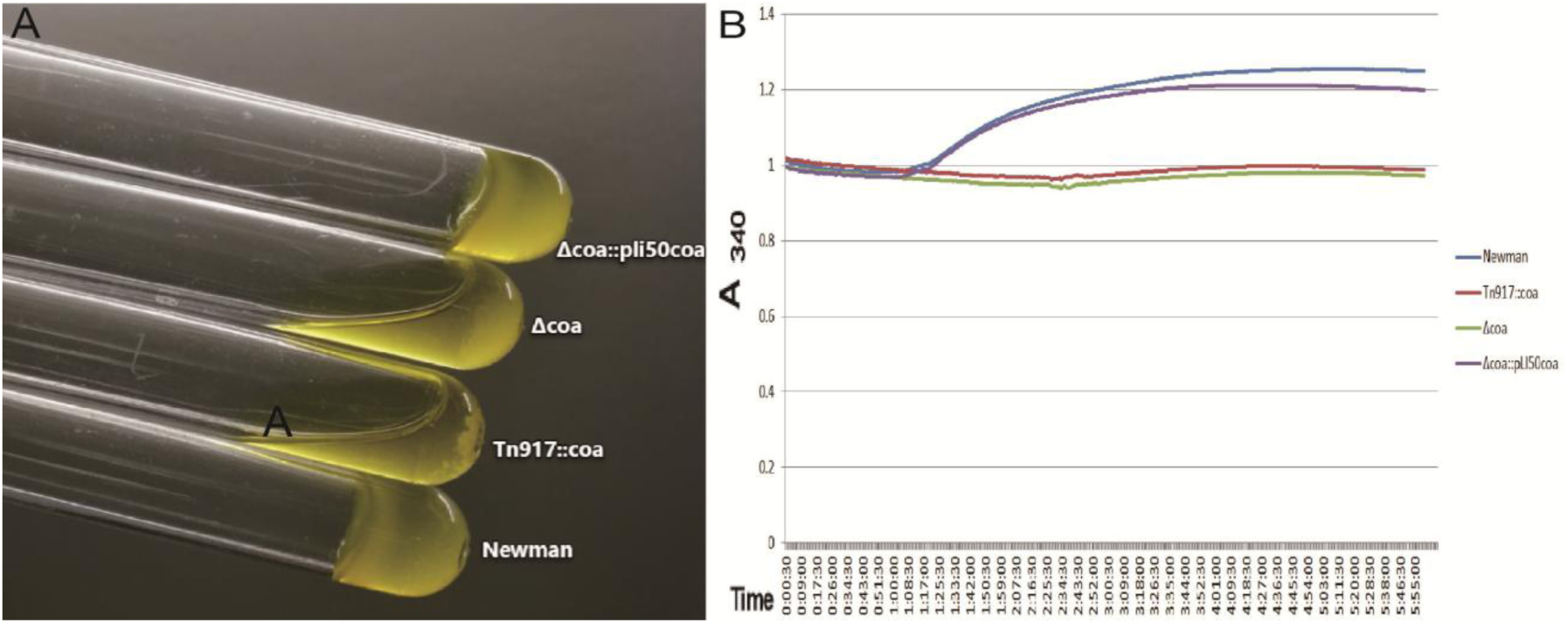
*Coa* is necessary to plasma coagulation. *Coa* gene was knockout, Newman had no plasma agglutinating ability; *coa* gene was complementation, Newman had recovery of plasma agglutination.

**Figure 2.**
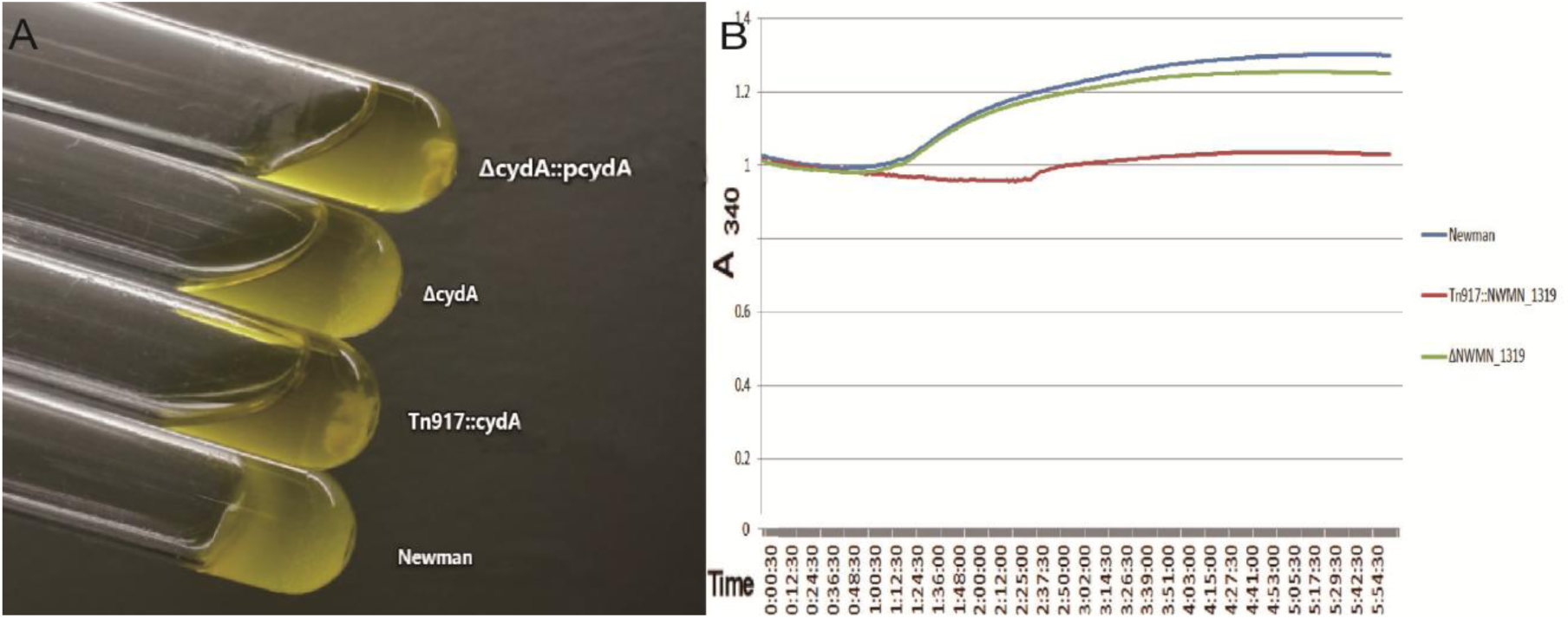
*cydA* is necessary to plasma coagulation. *cydA* gene was knockout, Newman had no plasma agglutinating ability; *cydA* gene was complementation, Newman had no recovery of plasma agglutination.

## References

1. Sirijatuphat R, Sripanidkulchai K, Boonyasiri A, Rattanaumpawan P, Supapueng O, Kiratisin P, et al. Implementation of global antimicrobial resistance surveillance system (GLASS) in patients with bacteremia. PLoS One, 2018; 13(1): e0190132

2. Buitron de la Vaga P, Tandon P, Qureshi W, Nasr Y, Jayaprakash R, Arshad S, Moreno D, et al. Simplified risk stratification criteria for identification of patients with MRSA bacteremia at low risk of infective endocarditis: implications for avoiding routine transesophageal echocardiography in MRSA bacteremia. Eur J Clin Microbiol Infect Dis, 2016; 35(2): 261–8

3. Monaco M, Pimentel de Araujo F, Cruciani M, Coccia EM, Pantosti A. Worldwide epidemiology and antibiotic resistance of Staphylococcus aureus. Curr Top Microbiol Immunol, 2017; 409: 21–56

4. Assis LM, Nedeljković M, Dessen A. New strategies for targeting and treatment of multi-drug resistant Staphylococcus aureus. Drug Resist Updat, 2017; 31: 1–14

5. Chen K, Lin S, Li P, Song Q, Luo D, Liu T, Zeng L, et al. Characterization of Staphylococcus aureus of Staphylococcus aureus isolated from patients with burns in a regional burn center, Southeastern China. BMC Infect Dis. 2018,18(1): 51

6. Tuchscherr L, Löffler B. Staphylococcus aureus dynamically adapts global regulators and virulence factor expression in the course from acute to chronic infection. Curr Genet, 2016; 62(1): 15–7

7. Hong X, Qin J, Li T, Dai Y, Wang Y, Liu Q, He L, et al. Staphylococcus aureus A promotes colonization and immune evasion of the epidemic healthcare-associated MRSA ST-239. Front Microbiol, 2016; 7: 951

8. Isobe H, Miyasaka D, Ito T, Takano T, Nishiyama, Iwao Y, et al. Recurrence of pelvic from panton-valentine leukocidin-positive community-acquired ST30 methicillin-resistant Staphylococcus aureus. Pediatr Int. 2013; 55(1): 120–3

9. Karauzum H, Adhikari RP, Sarwar J, et al. Structurally designed attenuated subunit vaccine for S.aureus LukS-PV and LukF-PV confer protection in a mouse bacteremia model. Plos One, 2013; 8(6): e65384.

10. McAdow M, DeDent AC, Emolo C, et al. Coagulases as determinants of protective immune responses against *Staphylococcus aureus* [J]. Infect Immun, 2012;80(10): 3389–98

11. Li M, Du X, Villaruz AE, et al. MRSA epidemic linked to a quickly spreading colonization and virulence determinant [J]. Nat Med; 18(5): 816–9

12. Hook JL, Islam MN, Parker D, Prince AS, Bhattacharya S, Bhattacharya J. Disruption of staphylococcal aggregation protects against lethal lung injury [J]. J Clin Invest. 2018; 128(3): 1074–1086

13. Peetermans M, Verhamme P, Vanassche T. Coagulase activity by Staphylococcus aureus: A potential target for therapy? Semin Thromb Hemost. 2015; 41(4): 433–44

14. Friedrich R, Panizzi P, Fuentes-Prior P, Richter K, et al. Staphylocoagulase is a prototype for the mechanism of cofactor-induced zymogen activation [J]. Nature, 2003; 425: 535–539.

15. McDevitt D, Nanavaty T, House-Pompeo K, Bell E, et al. Characterization of the interaction between the *Staphylococcus aureus* clumping factor (ClfA) and fibrinogen [J]. Eur J Biochem, 1997; 247: 416–424

16. Cheng AG, McAdow M, Kim HK, et al. Contribution of coagulases towards *Staphylococcus aureus* disease and protective immunity [J]. PLoS Pathog, 2010; 6: e1001036.

17. Josefsson E, Harford O, O’Brien L, et al. Protection against experimental *Staphylococcus aureus* arthritis by vaccination with clumping factor A, a novel virulence determinant [J]. J Infect Dis, 2001; 184: 1572–1580.

18. Weems JR, Domanski PJ, Patel PR, et al. Phase II, randomized, double-blind, multicenter study comparing the safety and pharmacokinetics of Tefibazumab to placebo for treatment of *Staphylococcus aureus* bacteremia [J]. Antimicrob Agents Chemother, 2006; 50: 2751–2755.

19. Sirichoat A, Wongthong S, Kanyota R, et al. Phenotypic characterizatics of vancomycin-non-susceptible *Staphylococcus aureus* [J]. Jundishapur J Microbiol, 2016;9(1): e26069.

20. Bae, T., and Schneewind, O. (2006). Allelic replacement in *Staphylococcus aureus* with inducible counter-selection. Plasmid 55, 58–63.

21. Wang W, Chen J, Chen G, Du X, Cui P, Wu J, et al. Transposon mutagenesis identifies novel associated with Staphylococcus aureus persister formation. Front Microbiol. 2015; 6: 1437

22. Loof TG, Goldmann O, Naudin C, Mörgelin M, Neumann Y, Pils MC, et al. Staphylococcus aureus-induced clotting of plasma is an immune evasion mechanism for persistence within the fibrin network. Microbiology. 2015; 161(Pt 3): 621–627

23. Kawada-Matsuo M, Oogai Y, Komatsuzawa H. Sugar allocation to metabolic pathways is tightly regulated and affects the virulence of Stretococcus mutans. Gene (Basel). 2016; 8(1): pii: E11

24. Bowman JP, Bittencourt CR, Ross T. Differential gene expression of Listeria monocytogenes during high hydrostatic pressure processing. Microbiology. 2008, 154(Pt): 462–75

25. Guo H, Hall JW, Yang J, Ji Y. The SaeRS two-component system controls survival of Staphylococcus aureus in human blood through regulation of coagulase. Front Cell Infect Microbiol. 2017; 7: 204

26. Flick MJ, Du X, Prasad JM, Raghu H, Palumbo JS, Smeds E, et al. Genetic elimination of the binding motif on fibrinogen for the S.aureus virulence factor ClfA improves host survival in septicemia. Blood, 2003; 121(10): 1783–94

27. Tennifer NW, Heidi AC, Adam RS, et al. The *Staphylococcus aureus* ArIRS two-component system is a novel regulator of agglutination and pathogenesis [J]. Plos Pathog, 2013; 9(12): e1003819.

28. Xue T, You Y, Shang F, Sun B. Rot and agr system modulate fibrinogen-binding ability mainly by regulating clfB expression in Staphylococcus aureus NCTC8325. Med Microbiol Immunol, 2012, 201(1): 81–92

29. Walker JN, Crosby HA, Spaulding AR, Salgado-Pabón W, Malone CL, Rosenthal CB, et al. The Staphylococcus aureus ArIRS two-component system is a novel regulator of agglutination and pathogenesis. PLoS Pathog, 2013; 9(12): e1003819

